# Integration of embryo-endosperm interaction into a holistic and dynamic picture of seed development using a rice mutant with notched-belly grains

**DOI:** 10.1101/2021.04.29.441907

**Authors:** Yang Tao, Lu An, Feng Xiao, Ganghua Li, Yanfeng Ding, Matthew J. Paul, Zhenghui Liu

## Abstract

The interaction between the embryo and endosperm affects seed development, an essential process in yield formation in crops such as rice. Signals that communicate between embryo and endosperm are largely unknown. Here we use the notched-belly (NB) mutant with impaired communication between embryo and endosperm to evaluate 1) the impact of embryo on developmental staging of the endosperm; 2) signaling pathways emanating from the embryo that regulate endosperm development. Hierachical clustering of mRNA datasets from embryo and endosperm samples collected through development in NB and wild type showed a delaying effect of the embryo on the developmental transition of the endosperm by extending the middle stage. K-means clustering further identified coexpression modules of gene sets specific for embryo and endosperm development. Combined gene expression and biochemical analysis showed that T6P-SnRK1, gibberellin and auxin signalling from the embryo regulate endosperm developmental transition. The data enable us to propose a new seed developmental staging system for rice and the most detailed signature of rice grain formation to date, that will direct genetic strategies for rice yield improvement.

## INTRODUCTION

Rice is a main contributor of dietary calories and nutrients for nearly half of the global population (Wu et al., 2020). Rice grain yield has to be sustainably improved both in quantity and quality, to solve formidable challenges including the ever-increasing population, climate change, and the quest for high-quality rice that has accompanied the rise in living standards (Sreenivasulu et al., 2015). A detailed understanding of the molecular and physiological mechanisms underpinning seed development may enable the design of effective strategies to boost rice quality and yield.

The seed (grain) of rice (*Oryza sativa*) is a complex but delicate biological system, containing three genetically distinct tissues, the diploid embryo, the triploid endosperm, and the diploid maternal tissues (An et al., 2020). Concomitant development of embryo and endosperm under the constraint of maternal tissues requires coordination of the two compartments (Lafon-Placette and Köhler, 2014; Doll et al., 2020). Growing evidence in rice as well as Arabidopsis (*Arabidopsis thaliana*) and maize (*Zea mays*) support bidirectional communication between embryo and endosperm. In brief, the endosperm plays crucial role in nurturing the developing embryo, whereas the embryo exerts a “counter-acting regulation force” on endosperm, as reviewed by An et al. (2020) and Ingram (2020).

Transcriptome analyses of seed development have already identified most genes expressed in its different compartments (Olsen, 2020). These comprehensive studies provided insights into the molecular networks and pathway interactions that function during the development of individual seed compartments for Arabidopsis (Belmonte et al., 2013), maize (Chen et al., 2014; Yi et al., 2019), wheat (*Triticum aestivum*; Xiang et al., 2019), and barley (*Hordeum vulgare*; Bian et al., 2019). In rice, laser-capture microdissection (LCM) has been employed to unravel the molecular mechanisms underpinning grain development and its response to abiotic stress (Ishimaru et al., 2019; Ram et al., 2020; Wu et al., 2020). Regardless of a large number of comprehensive studies on the transcriptome atlas for the developing seed with high spatio-temporal resolutions, detailed information about the communications between embryo and endosperm is still lacking, which is partly attributed to the complexity of seed structures (Doll et al., 2020).

Notched-belly grain is a misshapen type of rice grain, having a notched-line on its ventral side and thus being inferior in appearance and milling quality (Tong et al., 2018). Previously, we identified a notched-belly mutant (NB) of a japonica rice Wuyujing 3 using a chemical mutagen (Lin et al., 2014). It has high occurrence of white-belly grains, and the notched line is visible at 5 DAF, which separates the endosperm into two sub-compartments, the translucent upper part and the chalky bottom (Fig. 1). Since the formation of chalky tissue is a result of incomplete accumulation of starch and protein, it is tempting to speculate that the embryo plays a role in it, probably by depriving the nutrients from its proximate bottom endosperm (An et al., 2020). As revealed by our previous studies, the embryo demonstrated a negative effect on the contents of proteins, amino acids, and minerals in the chalky endosperm (Lin et al., 2016; Lin et al., 2017). Therefore, the NB mutant enables an integrated understanding of the cellular processes for rice grain formation from perspective of embryo-endosperm interaction.

**Fig. 1.**
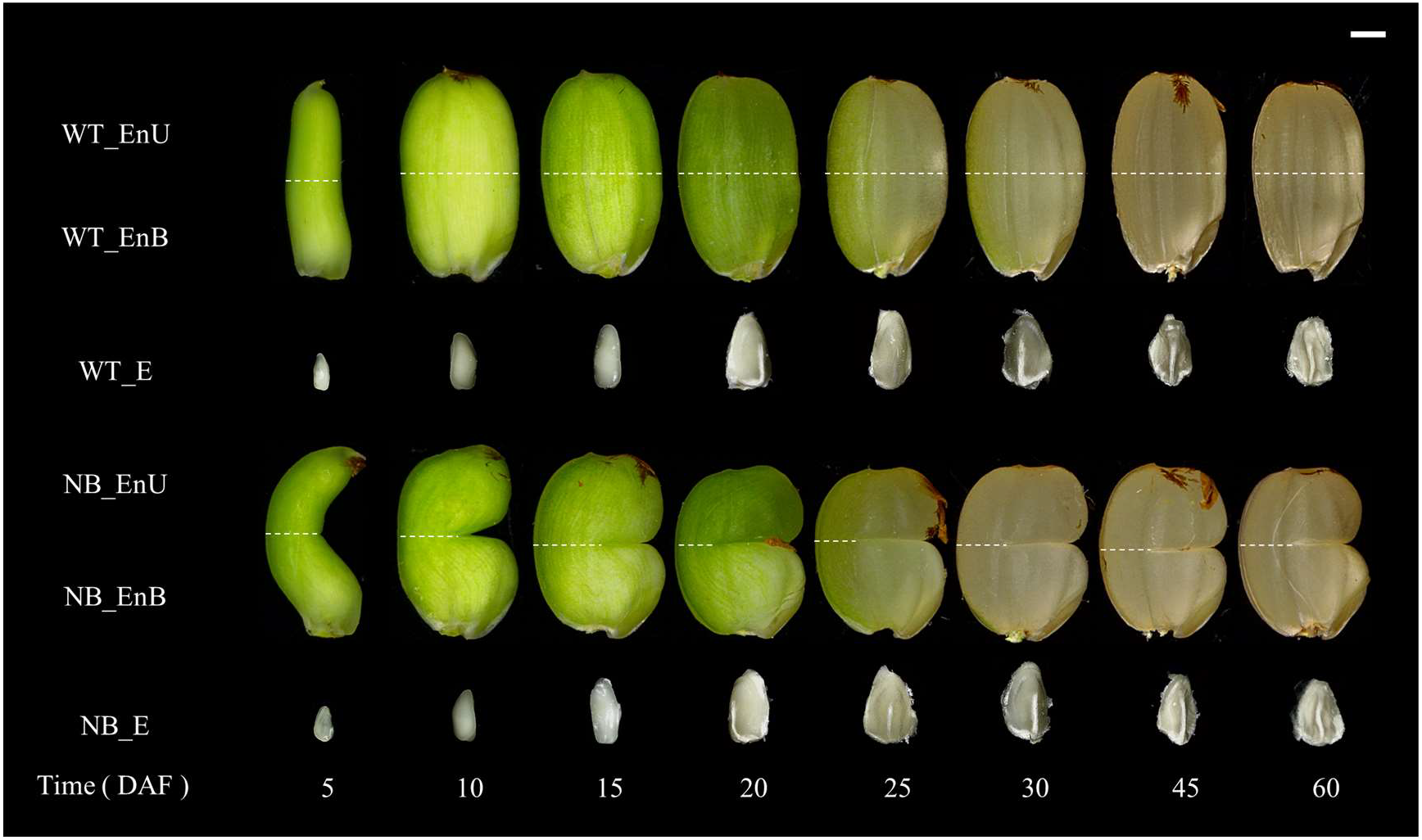
Overview of the time series samples of embryo and endosperms. Dotted line indicates the manual dissection of endosperm, cutting it into upper and bottom endosperms. DAF, days after fertilization; E, embryo; En, endosperm; EnB, bottom part of endosperm; EnU, upper part of endosperm; NB, notched-belly rice mutant; WT, wild type of Wuyujing-3. Bar = 1 mm

In this study, we used the NB mutant to characterize the genome-wide gene expression profile of embryo and endosperm samples between 0-60 DAF. In combination with physiological and histological investigations, we aimed to: (i) manifest the effect of embryo on endosperm development; and (ii) frame a holistic and dynamic picture of seed development by integrating the information of embryo-endosperm crosstalk. Comprehensive comparison of mRNAs datasets revealed a dragging effect of embryo on the developmental transition of the endosperm, which may be mediated by hormonal (GA and IAA) and T6P-SnRK1 signaling pathways. A new staging system is subsequently proposed by integrating the information about embryo-endosperm bidirectional communications. The findings obtained should offer insights into the biological processes and signatures of grain formation, hence providing a valuable resource for the manipulation of molecular and physiological processes responsible for cereal grain yield and quality.

## MATERIALS AND METHODS

### Plant materials and sampling

The notched-belly mutant (NB) was obtained by EMS treatment of a japonica rice cultivar Wuyujing3 (WT). It possesses a high ratio of notched-belly grains with a white-belly mainly on the bottom endosperm (Lin et al., 2014). In 2018, six seedlings of WT and NB were transplanted into a plastic pot filled with 10 kg paddy soil. The plants were grown under natural conditions, and at two days before flowering were transferred to a 31□/24□ growth chamber (12h/12h day and night cycle), with light intensity of 600 μmol photons m^−2^ s^−1^ and relative humidity 70 ± 5%. Flowering dates were carefully marked for the sampled caryopsis on the middle primary rachis. Grains were sampled at eight times, i.e. 5, 10, 15, 20, 25, 30, 45 and 60 DAF, with three biological replicates. Samples were quickly frozen in liquid nitrogen and then stored at −80 □ until analysis. The developing grains were manually dissected into three subsamples, the embryo, and the upper and bottom part of the endosperm (Fig. 1). It is noteworthy that the endosperm samples contain the maternal tissues such as pericarp and seed coat.

### Gene expression profiling using RNA-Seq

About 0.1 g sample was used for total RNA extraction by TRIzol reagent (Invitrogen, USA). The extracted RNA was dissolved in 100 μl of RNase-free water, and then quantified using a NanoDrop spectrophotometer (Thermo Scientific, USA). RNA quality was evaluated using the 6000 Pico LabChip of the Agilent 2100 Bioanalyzer (Agilent, USA). The qualification and quantification of the sample library were performed using an Agilent 2100 Bioanaylzer and StepOnePlus Real-Time PCR System (Applied Biosystem, USA). The library products were sequenced with BGISEQ-500 (BGI, China). Nipponbare reference genome (IRGSP-1.0; https://www.ncbi.nlm.nih.gov) was used to process the RNA-Seq libraries into read mapping and analysis.

To ensure good quality and effective mapping, the low-quality reads were removed using the SOAPnuke (V1.4.0) and trimmomatic (V0.36) software (Chen et al., 2018b), and only clean reads were processed for mapping using HISAT2 (V2.1.0) (Kim et al., 2015). The gene expression level was measured by RSEM (V1.2.8) software (Li and Dewey, 2011), and fragments per kilobase of transcript per million mapped reads (FPKM) values were used for quantifying gene activity. We only considered a gene as expressed if its FKPM value ≥ 1 to inhibit the influence of transcriptomic noise. To confirm the accuracy and authenticity of the three biological repeats, we calculated the Pearson correlation coefficient among them with the normalized expression level of log_2_ (FPKM value +1) (Supplementary Fig. S1). To identify differentially expressed genes (DEGs), we considered a DEG when it exhibits an absolute value of log_2_ ratio ≥1 compared with an FDR corrected P-value of ≤0.001 (Wang et al., 2010).

### Validation of transcriptome data using qRT-PCR

RNA-Seq data quality was validated by measuring the relative expression pattern of 12 genes functioning in C and N metabolism using qRT-PCR (Supplementary Fig. S2). Briefly, target-specific qRT-PCR primers were designed using the Primer5.0 software and synthesized, as presented in Supplementary Table S1. Each sample was represented by three biological and three technical replicates. The comparison of qRT-PCR results with RNA-Seq revealed the similar expression pattern between two datasets, thus validating the transcriptomic data (Supplementary Fig. S2). In addition, transcript abundance pattern of embryo (*OSH1* and *OsLEC1*) and endosperm *(GluD-1* and *RPBF)* specific genes also confirmed the credibility of the gene expression data (Supplementary Fig. S3), indicating that the embryo and endosperm samples were processed well.

### PCA and hierarchical clustering

To facilitate graphical interpretation of relatedness among the 48 tissue samples, we reduced the dimensional expression data to two dimensions by PCA through Omicshare tools (https://www.omicshare.com/tools/Home/Soft/pca). Hierarchical clustering (HCL) was carried out through Omicshare tools (https://www.omicshare.com/tools/Home/Soft/hca).

### Gene coexpression and functional enrichment analysis

The coexpression analysis for genes in embryo and endosperm was conducted using MeV (Howe et al., 2010). Notably, we used the Z-score value to calculate the relative expression levels before running the MeV for a tissue. These Z-score values were used as input for MeV and each individual dataset was clustered using the k-means method and Pearson correlation coefficients among the genes. Through Gene Ontology (GO) annotation (Ashburner et al., 2000), genes were annotated into the functional categories. Functional enrichment analysis was performed using the phyper function in R software with default setting (R Team, 2013). P-value was used as a filter mode, and significant GO categories (P-value <0.01) were identified. In addition, embryo or endosperm specific genes were identified by Z-score value. A gene was declared as embryo or endosperm specific if it had a Z-score above 2 in at least one of the embryo or endosperm samples as compared to all the same kind samples.

### Light microscopy observation of grain structure

Grains samples were fixed immediately in 0.1 M phosphate buffer solution (pH 7.2), containing 2.5% glutaraldehyde (v/v) and 2% paraformaldehyde (w/v). Afterwards, they were dehydrated in a graded ethanol series, transferred to acetone, and embedded in low-glutinosity Spurr’s resin (Spurr Low-Viscosity Embedding Kit, Sigma-Aldrich, St. Louis, USA). The sample blocks were polymerized at 70°C for 8 h. Semi-thin sections (2 μm) were cut at the longitudinal plane (embryo) and the transversal plane (endosperm) of the grains, using a Leica Ultrathin Microtome (EM UC7, Germany). Samples were stained with 1 % methyl violet, observed and photographed under a BX53 light microscope (Olympus, Tokyo, Japan).

### Chemical analysis for metabolites

#### Sucrose, fructose, glucose, and starch

Sugar contents were measured by ultra-high performance liquid chromatography system (UHPLC) as described by Chen et al. (2018a). Sucrose and hexoses were extracted with 80% (v/v) ethanol for 30 min at 80 °C, followed by centrifugation at 5000g for 15 min. The supernatant was pooled and lyophilized using a Maxi-Dry Lyo instrument (Heto-Holten, France), and resuspended in ultrapure water and filtered through a 0.45 μm pore size filter. The resulting filtrate was loaded into a Dionex Ultimate 3000 chromatography system. Starch content was measured using anthrone reagent at wavelength of 620 nm, after extraction by ethanol and digestion by perchloric acid.

#### T6P and SnRK1 activity

T6P was extracted and purified according to the method of Delatte et al. (2009), and quantified by ultra-performance liquid chromatography (ACQUITY UPLC I-Class PLUS, Waters, USA) coupled with triple quadruple mass spectrometry system (Xevo TQ-S micro, Waters, USA), following the method of Sastre Toraño et al. (2012). SnRK1 was extracted and its activity was assayed by the method of Zhang et al. (2009), using AMARA peptide as substrates.

#### Amino acids and proteins

Samples were boiled with 80% ethanol until totally bleached to extract free amino acids (FAAs). Then, the total volume was kept constant with distilled water and was centrifuged for 15 min. The supernatant was pooled and filtered through a 0.45 μm pore size filter, and then used for FAAs determination by amino acid analyzer (L-8900, Hitachi, Japan). Protein fractions including albumin, globulin, prolamin and glutelin were extracted and measured by the methods of Ning et al. (2010).

#### Phytohormones and minerals

IAA, ABA, CKs (tZ and tZR) and GAs (GA_1_, GA_3_, GA_4_, and GA_7_) were extracted and purified according to the method of Wu et al. (2016a). IAA, ABA and CKs were quantified by ultra-performance liquid chromatography (ACQUITY UPLC H-CLASS, Waters, USA) according to Chou et al. (2000). GAs were quantified by ultra-high performance liquid chromatography (AGLIENT1290, Agilent Technologies, USA) coupled with mass spectrometry system (SCIEX-6500Qtrap, AB SCIEX, USA). Minerals, including P, K, Ca, Na, Mg, Fe, Zn, Mn, and Cu, were quantified by Inductively Coupled Plasma Optical Emission Spectrometry (ICP-OES 710, Agilent Technologies, USA), according to the method of Wang et al. (2018).

### Statistical analysis

The data of metabolites presented in the study are averages of triplicate observations. Statistical analysis was performed with SPSS (V19.0, IBM) statistical software. Multiple comparisons were operated by Duncan’s multiple range test (P < 0.05). Origin 9.0 was used for plotting.

## RESULTS

### The generation and analysis of transcriptome data of developing embryo and endosperm

To systematically investigate the dynamics of transcriptome over seed development, we generated RNA-Seq libraries of three different seed tissues of the wild type WYJ3 (WT) and its notched-belly mutant (NB) from 8 different time points, including 16 embryos, 16 upper endosperms and 16 bottom endosperms (Fig. 1). In total, we detected 20820 genes expressed in at least one of the 48 samples. Among them, 18171 genes were expressed in both embryo and endosperm, while only 1969 and 680 genes were specifically expressed in embryo and endosperm, respectively (Fig. 2A). Subsequently, mRNA datasets from different tissues were compared to get a more specific look into the individual characteristics of the embryo and endosperm during seed development. Results showed that most of genes were expressed in seed tissues during the early developmental phase (Fig. 2B). Within endosperm, both genotypes showed higher gene activity in upper part of endosperm (EnU) than the bottom part (EnB). In addition, embryo samples had higher number of expressed genes as compared to those of the endosperm throughout seed development (Fig. 2C).

**Fig. 2.**
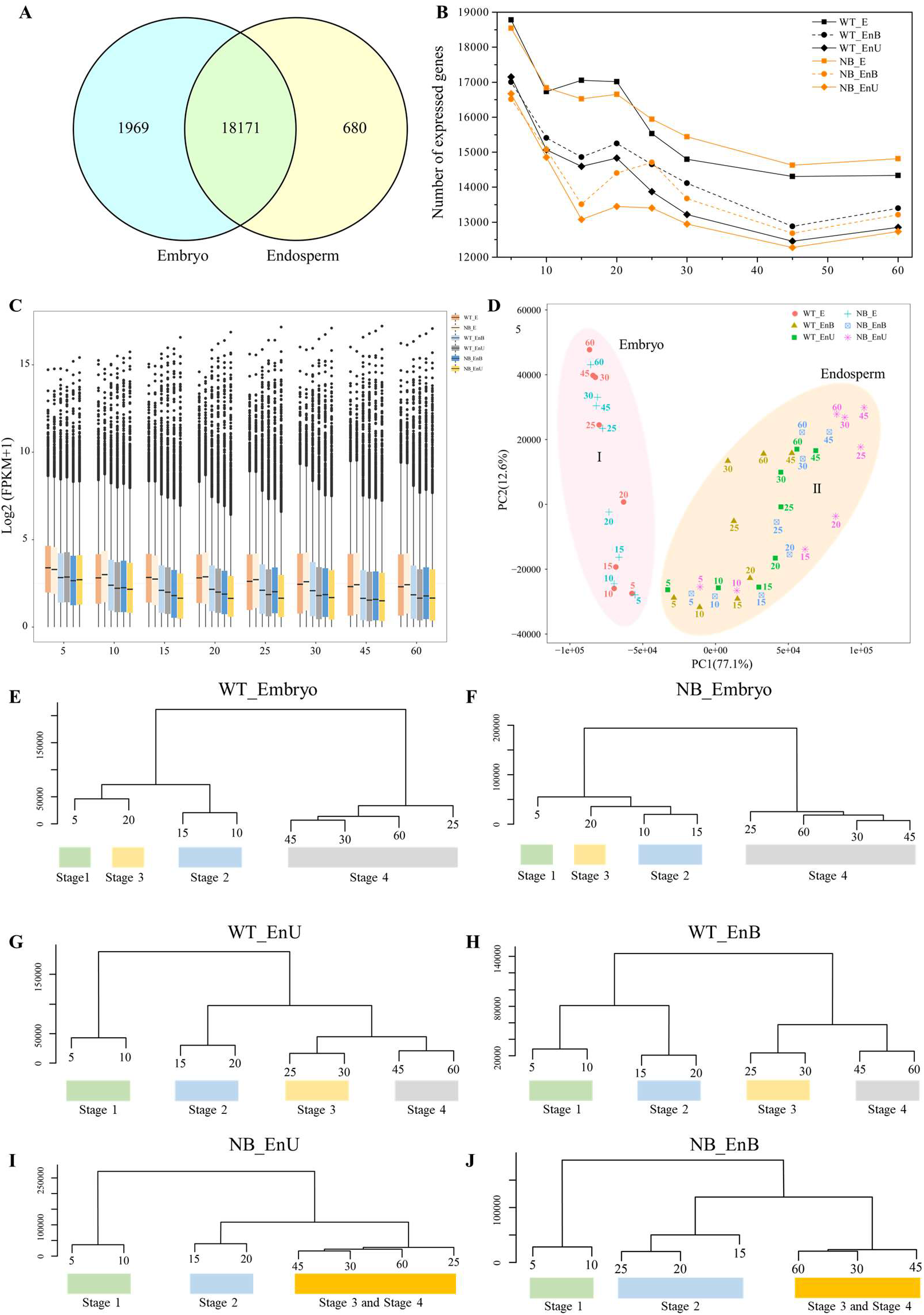
Global transcriptome relationships among tissues and developmental stages. (A) Venn diagram of the 20820 genes expressed in the embryo and endosperm. (B) Number of genes detected in each tissue. (C) Comparison of gene activity between embryo and endosperm. (D) PCA of the seed tissue mRNA populations. PCA plot shows two distinct groups of embryo and endosperm mRNA populations: group I for embryo and group II for endosperm. (E) and (F) Clustered dendrogram showing global transcriptome relationships of time series samples from the embryo of WT and NB, respectively. (G) to (H) Clustered dendrogram showing global transcriptome relationships of the upper and bottom endosperms of WT, respectively. (I) to (J) Clustered dendrogram showing global transcriptome relationships of the upper and bottom endosperms of NB, respectively. The bottom row indicates the developmental phases according to the cluster dendrogram of the time series data. Except in (A), data of 5, 10, 15…, and 60 represent sampling timepoints (days after fertilization).

### Categorizing developmental processes of embryo and endosperm based on gene expression pattern

PCA distinguished the samples into two distinct groups based on tissue identity, validating the purity of the tissue samples (Fig. 2D). Particularly, the first component (PC1; 77.1% variance explained) clearly separated the embryo from endosperm samples, while the second component (PC2; 12.6% variance explained) discriminated the developmental stages. We further classified the transcriptomic data into a dendrogram by hierarchical clustering (HCL) analysis, and uncovered four distinguishing groups within embryo and endosperm, with each group corresponding to a specific developmental stage (Fig. 2E-H).

For embryo samples, both genotypes showed an almost identical dendrogram (Fig. 2E, F). The first cluster is formed at 5 DAF, representing the stage around differentiation (Em-S1). During this stage, coleoptile and SAM emerge and first leaf primordium becomes visible on the opposite side of the coleoptiles, as visualized by electron microscopy (Supplementary Fig. S4). The second cluster from 10-15 DAF represents the embryo enlargement stage (Em-S2), as characterized by the formation of second and third leaf primordia, and organ enlargement (particularly the scutellum). The third cluster at 20DAF corresponds to the maturation phase (Em-S3) that shows no obviously morphological change. The fourth cluster from 25 DAF to 60 DAF represents the period of seed dormancy (Em-S4).

By contrast, the duration of developmental stages in endosperm varied with genotypes. Firstly, endosperms from WT showed four primary clusters (Fig. 2G, H), but those from NB showed only three clusters, suggesting the abnormal development during middle and later stages (Fig. 2I, J). Secondly, within the endosperm, the upper and bottom parts of WT showed identical duration. However, for NB, the time span of endosperm filling in the bottom endosperm (NB_EnB) was prolonged (15-25 DAF), as compared to the upper endosperm (NB-EnU) that had the identical duration of second cluster (15-20 DAF) to the WT.

Consequently, the maturation and dormancy stage were delayed in the bottom endosperm and displayed as a merged cluster (30-60 DAF) of stages 3 and 4, as compared to the upper part, where stages 3 and 4 cluster had the same duration (25-60 DAF) as WT. Collectively, these findings indicate a disturbance in the developmental process of the bottom endosperm of NB by its proximity to the embryo.

Therefore, normal type of WT instead of the disturbed NB is used for a general description of the developmental signatures of rice grain. The four clearly defined clusters in WT endosperm show that the differentiation (En-S1) stage last from 5-10 DAF in which it completes the cellularization, with the aleurone cells starting to become morphologically distinct from the starchy endosperm (Supplementary Fig. S4). Then storage accumulation stage (En-S2) occurs at 15-20 DAF, followed by the maturation (En-S3) phase of the endosperm at 25-30 DAF. The biological processes involved in dormancy (En-S4) are induced from 45-60 DAF. Altogether, these results suggested a divergent staging mechanism of developmental characteristics between embryo and endosperm based on transcriptomic data.

### Coexpressed gene sets of embryo and endosperm development

Using transcript datasets of WT, tissue-specific genes were clustered into coexpression modules by the k-means clustering algorithm (Howe et al., 2010). Gene ontology (GO) annotation was performed to assign genes to functional categories for each module (Ashburner et al., 2000). Twenty-two coexpression modules were generated for both embryo and endosperm (Fig. 3A, B). Among these modules, 11 and 9 modules were expressed broadly at more than one stage in embryo and endosperm, respectively, indicating some common cellular processes across several stages. Moreover, genes from the 11 modules of embryo and the 13 modules of endosperm were more prevalent at one out of the four developmental stages, indicating specific function of these modules at the corresponding stage.

**Fig. 3.**
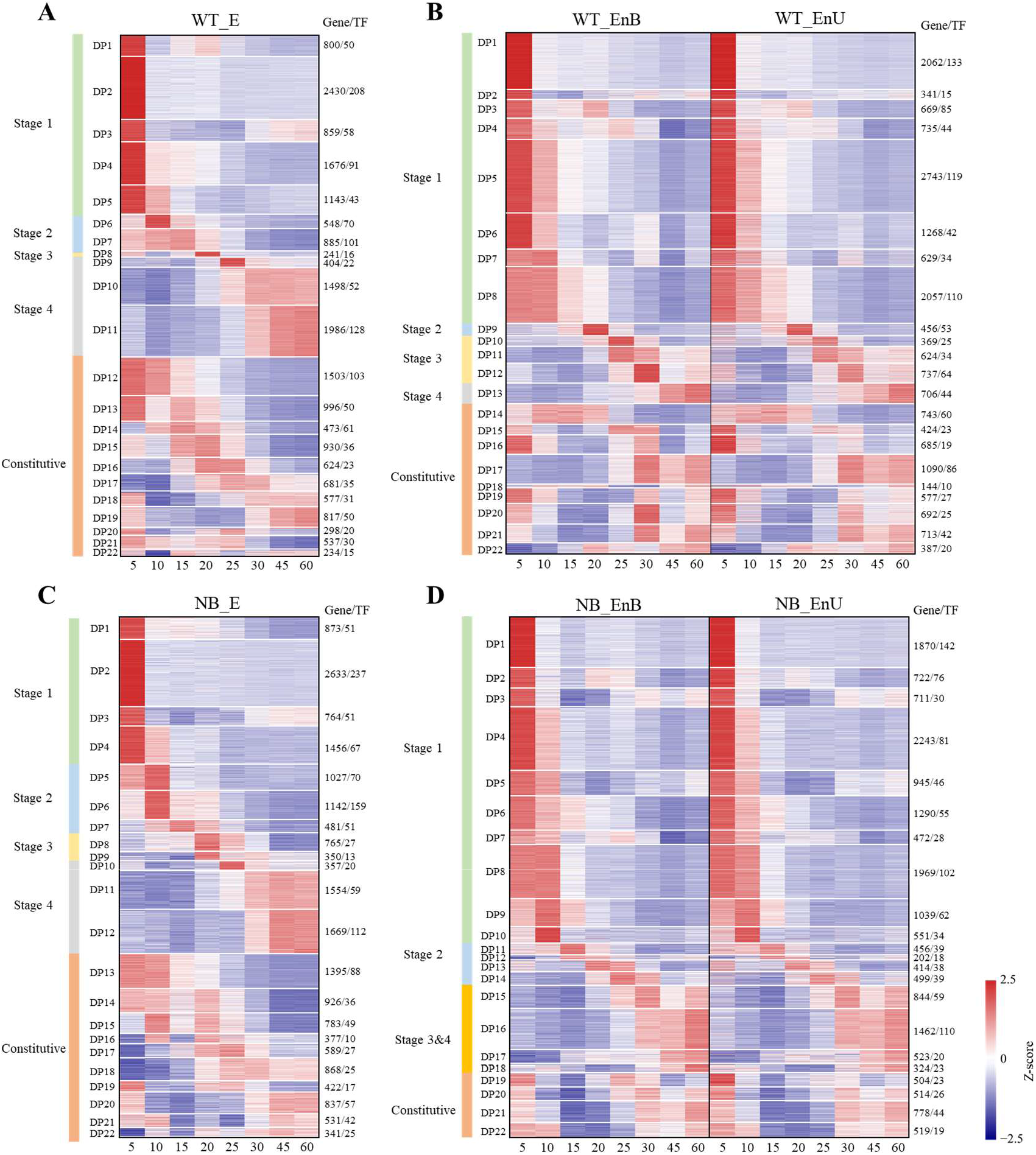
Expression patterns of genes in different coexpression modules for embryo and endosperm of WT (A, B) and NB (C, D). Coexpression modules are ordered according to the sample time points of their peak expression. For each gene, the FPKM value normalized by the maximum value of all FPKM values of the gene over all time points is shown. The numbers of genes and TFs in each module are shown on the right.

#### Cellular processes in the developing embryo

Cellular processes characterizing each developmental stage were identified by GO terms that were overrepresented at particular coexpression modules (Fig. 3A; Supplementary Fig. S5A). The early stage of Em-S1, best represented by modules DP1-DP5, was typified by the overrepresentation for GO term Ran GTPase binding, mitotic cell cycle, nuclear division, nuclear DNA replication, cell wall organization or biogenesis, cell differentiation, mitochondrial protein complex and tricarboxylic acid cycle. Em-S2 featured modules DP6 and DP7 that were enriched in GO term characteristic of embryo enlargement, including the cell division, cell wall organization or biogenesis and lipid droplet. The module DP8 presented an up-regulation of genes involved in nutrient reservoir activity and response to external biotic stimulus, suggesting the maturation of embryo characterized by Em-S3. Stage Em-S4, represented by modules DP9-DP11, was characterized by the overrepresentation of genes involved in response to biotic stimulus, structural constituent of ribosome, mitochondrial matrix, vacuole organization and photomorphogenesis. In addition, genes from modules DP12-DP22 expressed broadly in embryo across the sampled time points, which were related to protein complex, Golgi apparatus, protein kinase activity, cellular response to stimulus, RNA modification, ribosome biogenesis and endonuclease activity.

#### Cellular processes in the developing endosperm

The functional characterization of endosperm was found different from that of embryo as different number of modules presented a particular growth stage (Fig. 3B; Supplementary Fig. S5B). Modules DP1-DP8 were characterized as the stage of En-S1, including genes related to mitotic cell cycle, DNA replication, cell wall organization, tricarboxylic acid cycle and starch biosynthetic process. En-S2 featured the overrepresentation of GO term nutrient reservoir activity and transporter activity in DP9. Modules DP10-DP12 presented high expression of genes responsible for response to biotic stimulus and lipid storage, representing endosperm maturation at stage En-S3. Finally, the module DP13 presented an up-regulation of the genes involved in positive regulation of autophagy, organelle disassembly, metal ion transport, and cellular catabolic process, characteristic of the desiccation and dormancy stage of endosperm (En-S4). Notably, genes responsible for metal cluster binding, macromolecular complex, RNA processing and signal transducer activity were overrepresented in DP14-DP22 and were broadly expressed throughout development.

### Metabolic dynamics of embryo and endosperm development

#### Sugars and starch

During embryo and endosperm development, sucrose content gradually increased, peaking at 15 DAF and then decreased thereafter (Fig. 4A; Supplementary Fig. S6A). Glucose and fructose levels decreased during seed development (Supplementary Figs. S7A, S7B, S8A, S8B). The ratio of glucose to sucrose mediates endosperm differentiation (Olsen, 2020). It was higher in the embryo and endosperm at 5-10DAF, and then decreased gradually (Supplementary Figs. S7C, S8C), corresponding to the transition from differentiation to storage accumulation. These sugars were unevenly distributed in rice seed, with the embryo generally having higher contents (Fig. 4A; Supplementary Figs. S6A, S7A, S7B, S8A, S8B). Genes participating in sugar metabolism were differentially expressed between embryo and endosperm. For the five genes encoding sucrose synthase, two of them (*OsSUS1* and *OsSUS6*) were mainly expressed in embryo, while another three (*OsSUS2*, *OsSUS3*, *OsSUS4*) in endosperm (Fig. 4A; Supplementary Fig. S6A).

**Fig. 4.**
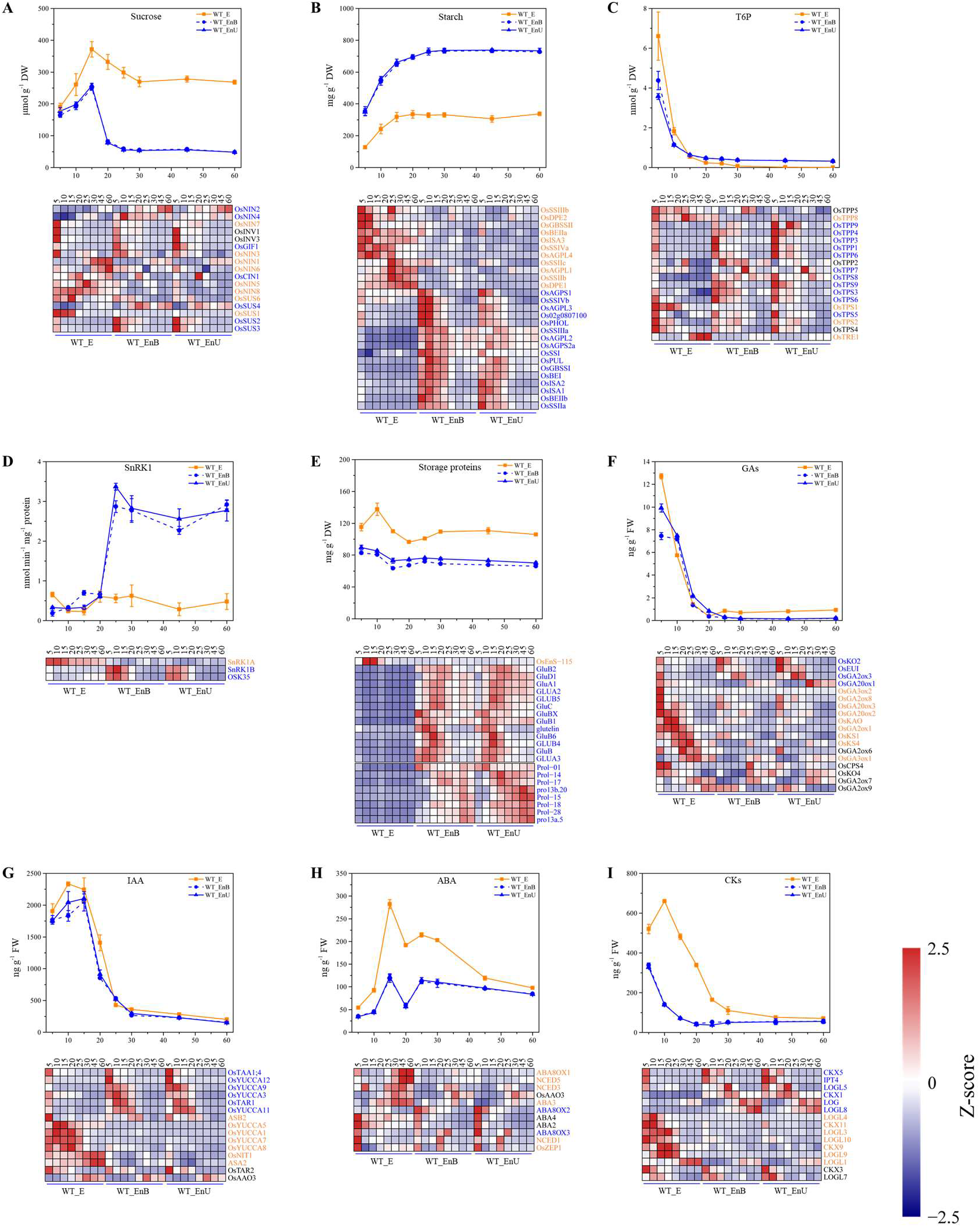
Carbohydrates, proteins, SnRK1, and hormones, and their regulating genes in the embryo and endosperm of WT throughout development. Each value represents the mean ± SE of three replicates. Black, orange, and blue indicate non-tissue, embryo, and endosperm specific genes, respectively.

During grain filling, starch content increased gradually, peaking at 20 DAF and 30 DAF in the embryo and endosperm, respectively (Fig. 4B; Supplementary Fig. S6B). Compared with embryo, endosperm contained higher starch content, which is consistent with previous report (Juliano and Bechtel, 1985). Starch synthesis-related genes were coordinately expressed in the embryo and endosperm, similar to that in wheat (Xiang et al., 2019). Moreover, these genes were primarily expressed in endosperm at 5-20 DAF. By contrast, they displayed a gradual increase in embryo at 5-20 DAF.

#### *T6P and its target protein* SnRK1

Trehalose-6-phospate (T6P) signals the availability of sucrose in plant cells through the feast-famine kinase, SnRK1 (O’Hara et al., 2013; Paul et al., 2017; Dingkuhn et al., 2020). T6P level in embryo and endosperm showed a gradual decrease in the development course. Compared with endosperm, embryo has higher content at early stage (5-10DAF) (Fig. 4C; Supplementary Fig. S6C). For example, at 5 DAF, T6P was 3.98 and 6.61 nmol g^−1^ DW in endosperm and embryo, respectively. In addition, *TPS* genes were expressed in both embryo and endosperm, being predominantly expressed at 5-20 DAF.

The SnRK1 activity in endosperm maintained a low level at 5-20 DAF, and then increased rapidly at later stages (25-60 DAF). Different from the endosperm, SnRK1 in the embryo was at a lower level, being higher only at 5 DAF (Fig. 4D; Supplementary Fig. S6D). *SnRK1A* was specifically expressed in the embryo, while two *SnRK1B* genes (*SnRK1B* and *OSK35*) were mainly expressed in endosperm. In addition, *SnRK1A* was broadly expressed across the time points in embryo. By contrast, *SnRK1B* and *OSK35* were more prevalent in endosperm at 5-20 DAF.

#### Amino acids and proteins

During embryo development, the level of total free amino acids (FAAs) gradually decreased at 5-25 DAF and increased thereafter (Supplementary Figs. S7D, S8D). In contrast, FAAs in endosperm decreased as grain filling progressed. In addition, FAAs were higher in embryo than endosperm. Storage proteins were higher in embryo than endosperm (Fig. 4E; Supplementary Fig. S6E). However, gene expression pattern of storage protein synthesis showed an opposite correlation with storage proteins content, suggesting that their encoding genes are specifically expressed in the endosperm, as in wheat (Xiang et al., 2019).

#### Phytohormones

GAs showed a gradual decrease over seed developmental course. Compared with endosperm, embryo had higher content at 5 DAF, while exhibited an opposite trend at 10 DAF (Fig. 4F; Supplementary Fig. S6F). GA-synthetic genes, *KS* (*OsKS1* and *OsKS4*), *GA20ox* (*OsGA20ox2* and *OsGA20ox3*), and *GA3ox* (*OsGA3ox1* and *OsGA3ox2*), were uniquely expressed in embryo at 5-20 DAF, implying that the embryo may be main site for GAs synthesis (Xue et al., 2012; Zhang et al., 2020a). In addition, two GA-deactivating genes, *EUI* and *OsGA2ox3*, were specifically enriched in endosperm at 5 DAF and 15-20 DAF, respectively.

Auxin gradually increased until 10-15 DAF in embryo and endosperm, then decreased thereafter (Fig. 4G; Supplementary Fig. S6G). Compared with endosperm, embryo had higher content at 5-20 DAF. Notably, *OsTAA1;4* was mainly expressed in endosperm at 5 DAF.

ABA content in endosperm was half of that in embryo. It showed two peaks at 15 DAF and 25 DAF in embryo and endosperm (Fig. 4H; Supplementary Fig. S6H), in line with the biphasic pattern of the embryonic development (Qin et al., 1990). Activity of ABA-synthesis genes in embryo and endosperm displayed two peaks. In embryo, *NCED1*, *OsZEP1* and *ABA2* were highly expressed at 5 DAF, while *NCED3* and *NCED5* at 25 DAF. In endosperm, *NCED1*, *OsZEP1* and *ABA2* were highly expressed at 5 DAF, while *NCED1* and *OsZEP1* at 20 DAF.

*OsIPT4* were specifically expressed in endosperm, especially at 5-10 DAF (Fig. 4I; Supplementary Fig. S6I). In embryo, CKs levels gradually increased, peaked at 10 DAF and then decreased thereafter, consistent with the expression patterns of CKs-synthetic genes. The CKs in endosperm was lower than those in embryo at 5-30 DAF and peaked at 5 DAF.

#### Minerals

In embryo, contents of K, Ca, Na, Fe, Mn, Cu and Zn decreased as development progressed, while those of Mg and P gradually increased (Fig. 5; Supplementary Fig. S9). In endosperm, contents of the nine mineral nutrients decreased as grain filling progressed, being higher at early stage than the later stage. This could be attributed to the dilution effect of grain dry matter, which had a higher rate of accumulation than the minerals (Wang et al., 2018). Compared with endosperm, contents of all minerals were high in the embryo.

**Fig. 5.**
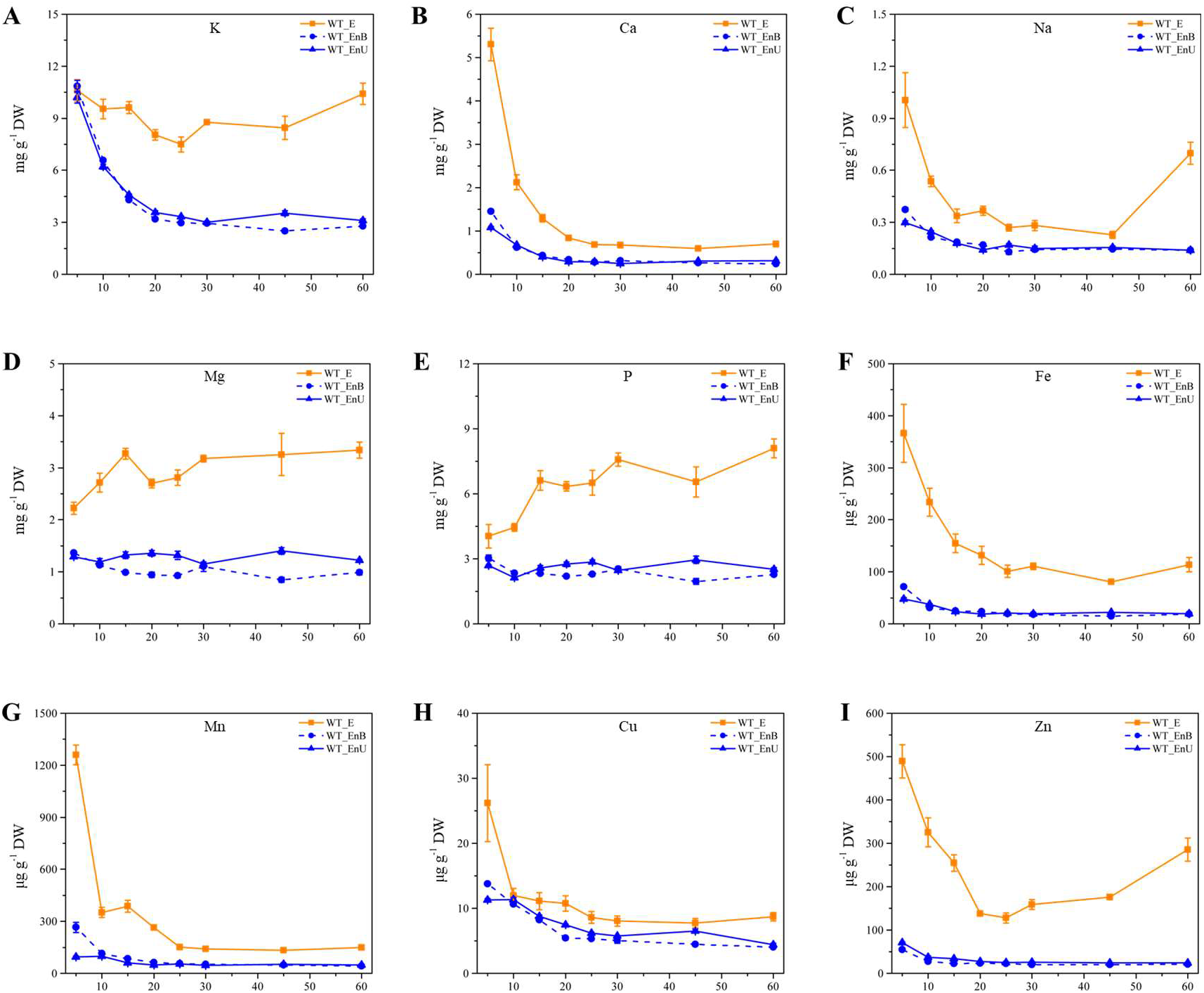
Minerals in the embryo and endosperm of WT and NB across the developmental stages. Each value represents the mean ± SE of three replicates.

### Influence of embryo on the transition of developmental stages of endosperm

#### A novel method to qualify the effect of embryo on endosperm development

As shown in Fig. 2, the duration of middle stage in the bottom part of the NB grain (Fig. 2J) was prolonged relative to that of its upper counterpart (Fig. 2I) as well as those of the WT (Fig. 2G, H), indicating a strong influence of the embryo on the bottom endosperm proximate to it. To quantitively evaluate the embryo effect, we developed a novel comparison system by comparing the upper and bottom endosperms of NB, using WT as a reference (Fig. 6). Briefly, this system has three key components: (i) Position effect. The comparison between the upper (WT_EnU) and bottom (WT_EnB) endosperms of WT reflects the difference in position between upper and bottom endosperms within the WT grain, where nutrients and signals move freely without being blocked by the notched line as in NB (Fig. 6A). (ii) Compound effect. Due to the notched line, movement of nutrients and signals are severely restricted between the two endosperms, trapping the influence of the embryo in the bottom endosperm. Comparison between the bottom part (NB_EnB) and upper part (NB_EnU) of the NB grain shows the compound effect of position and embryo (Fig. 6B). (iii) Embryo effect. Finally, by eliminating the position effect, we can precisely calculate the influence of embryo on the endosperm via NB_(EnB/EnU)_/ WT_(EnB/EnU)_ (Fig. 6C).

**Fig. 6.**
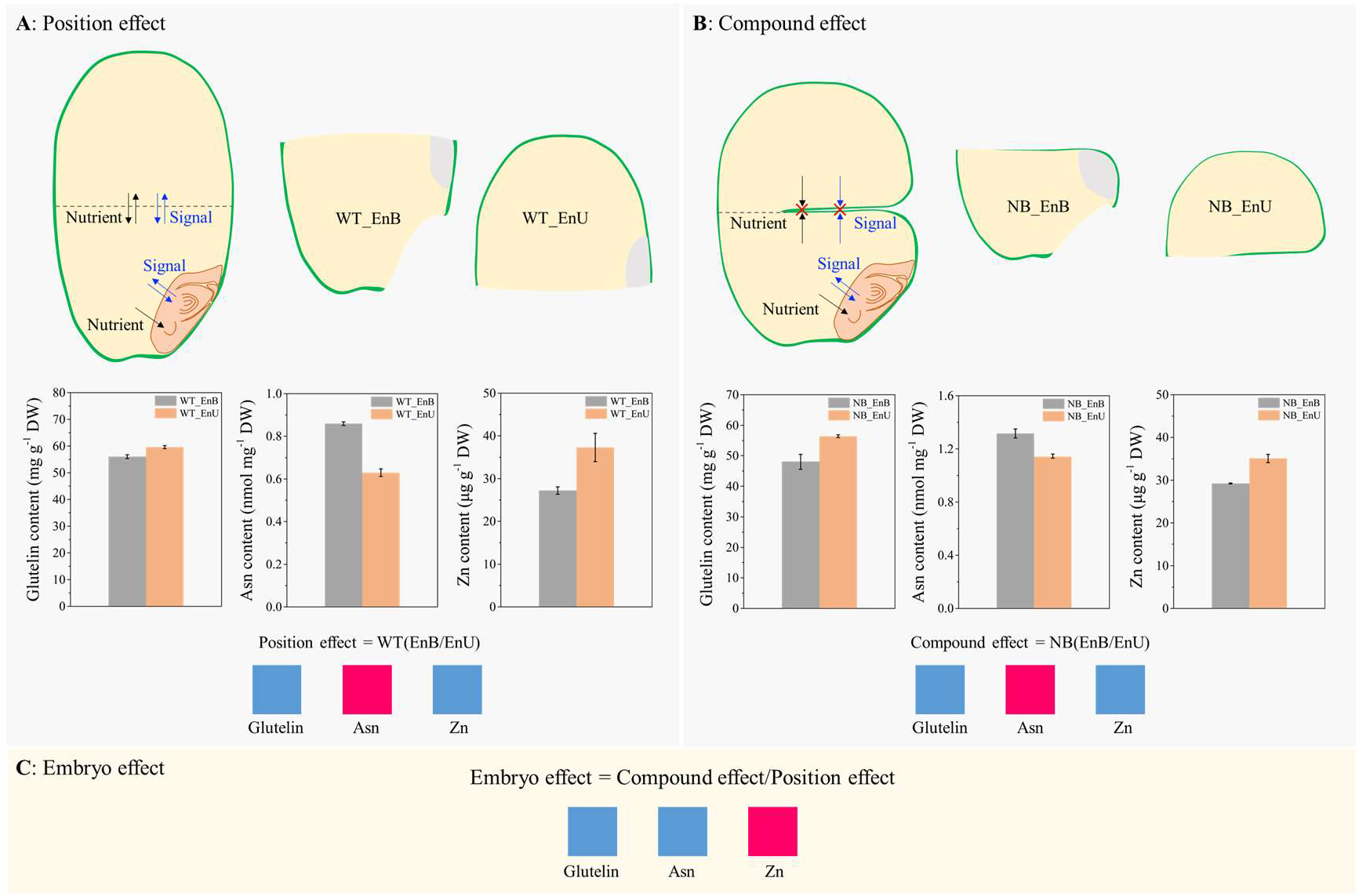
Diagrammatic representation of the new comparison method for quantifying the embryo effect on endosperm development. This method assembles three key components: (A) Quantification of the position effect (PE) by comparison between the upper (WT_EnU) and bottom (WT_EnB) endosperms of WT. This reflects the difference in position between upper and bottom endosperms within the WT grain, where nutrients and signals move freely without being blocked by the notched line as in NB. (B) Quantification of the compound effect (CE) of position and embryo by comparison between the upper (NB_EnU) and bottom (NB_EnB) endosperms of NB. Due to the notched line, movement of nutrients and signals are severely constrained between the two endosperms. Therefore, the influence of the embryo is trapped and thus mainly expressed in the bottom endosperm. Note that 2/3 interface between the upper and bottom is cut down by the notched line. (C) Evaluation of the embryo effect (EE) by secondary comparison of CE/PE to exclude the position effect. Three metabolites are used to exemplify the working principle of this method. For example, Asn content is up-regulated in the bottom endosperm of both WT and NB. By contrast, comparison of NB and WT reveals that the embryo effect is down-regulating it in the endosperm. Red and navy blue squares indicate up and down-regulation in the bottom endosperm, respectively.

To exemplify the working principle of this comparison method, we selected three metabolites including glutelin, asparagine (Asn), and Zn to show the influence of embryo (Fig. 6). Firstly, taking in account the position effect and compound effect, the up-regulation of Asn and down-regulation of glutelin and Zn was observed in both genotypes. However, when eliminating the effect of position, embryo has a positive effect on the content of Zn and a negative effect Asn and glutelin, verifying the influence of embryo on endosperm filling.

#### T6P-SnRK1 signaling putatively mediates the crosstalk between embryo and endosperm

Using the new comparison method, our results showed that T6P-SnRK1 signaling pathway was active in the endosperm, consistent with that of the wheat (Martínez-Barajas et al., 2011). As shown in Fig. 7, at early stage of 5 to 10 DAF, embryo had a negative effect on sucrose, glucose and fructose levels in endosperm during 5-10 DAF, probably due to its nutrient consumption. The decrease of sucrose content in endosperm was accompanied by the decline in T6P levels, which in return inhibited SnRK1 activity (Fig. 7A). SnRK1 promoted catabolic activity in the endosperm, as evidenced by the declining accumulation of starch and storage proteins (prolamin and glutelin), and the up-regulation of genes encoding amylase (*AMY3D*), lipase (*Os01g0651800*, *Os01g0710700*, and *Os05g0574100*), and protease (*OsSAG12* and *OsSCP28*) in endosperm. Consistently, genes participating in synthetic process of starch (*OsAGPL4*, *OsSSSIIIa*, *OsSSIVa*, *OsSSIVb*, and *Os02g0807100*) and storage protein (*OsEnS-115*, *Prol-14*, *Pro-15*, *Prol13b.20*, and *Glutelin type-B 2-like*) were all down-regulated at 5-10 DAF (Fig. 7B).

**Fig. 7.**
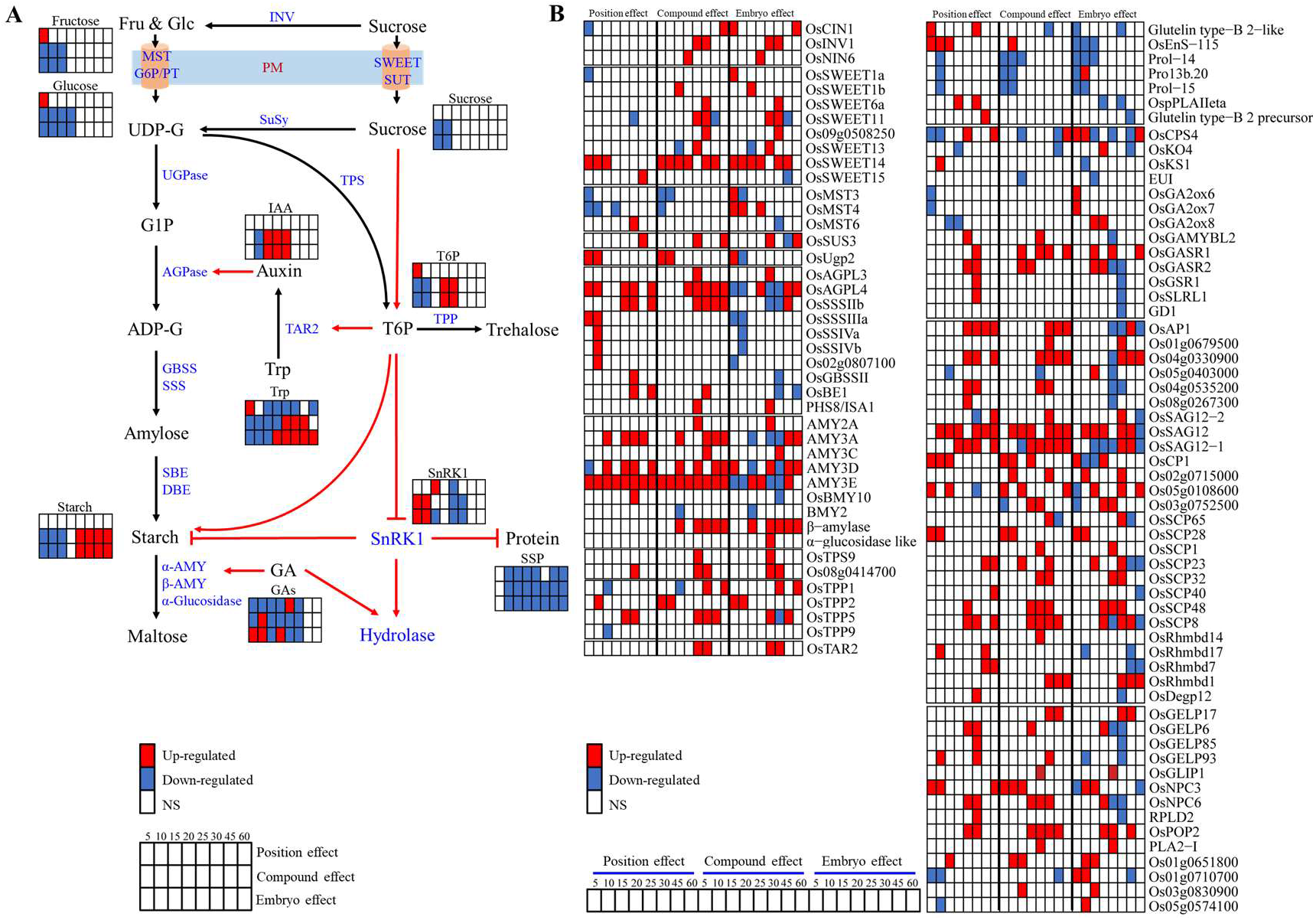
Putative role of T6P-SnRK1 signaling in developmental transition of the endosperm as revealed by the proposed new comparison method. Effects of position, compound, and embryo are all shown. (A) Schematic diagram showing the putative T6P-SnRK1 signaling pathway. (B) Heatmap of differentially expressed genes associated with position, compound, and embryo effects. ADP-G, ADP-glucose; AGPase, ADP-glucose pyrophosphorylase; DBE, debranching enzyme; Fru, fructose; G6P/PT, glucose-6-phosphate/phosphate translocator; G1P, glucose 1-phosphate; GA, gibberellin; GBSS, granule-bound starch synthase; Glc, glucose; IAA, indoleacetic acid; INV, invertase; ISA, isoamylase; MST, monosaccharide transporter; PM, plasm membrane; SBE, starch branching enzyme; SnRK1, sucrose non-fermenting-1 related protein kinase 1; SSP, seed storage protein; SSS, soluble starch synthase; SuSy, sucrose synthase; SUT, sucrose transporter; T6P, trehalose 6-phosphate; TAR2, tryptophan aminotransferase 2; TPP, trehalose 6-phosphate phosphatase; TPS, trehalose 6-phosphate synthase; Trp, tryptophan; UDP-G, UDP-glucose; UGPase, UDP-glucose pyrophosphorylase; α-AMY, α-amylase; β-AMY, β-Amylase. Red and navy rectangles indicate up- and down-regulation, respectively.

Conversely, at middle stage between 20-25 DAF, T6P-SnRK1 signaling showed an opposite trend relative to that between 5-10 DAF. Gene activities of sucrose metabolism (*OsINV1* and *OsNIN6*) and T6P synthesis (*OsTPS9* and *Os08g0414700*) were up-regulated (Fig. 7). SnRK1 activity was inhibited synchronously in endosperm (Fig. 7A). Meanwhile, starch synthetic process was promoted and catabolic processes were inhibited by SnRK1, respectively, as reflected by the overrepresented starch-synthetic genes (*OsAGPL3* and *ISA1*) and underrepresented catabolic genes encoding amylase (*AMY3A* and *AMY3E*), and protease (*OsAP1*, *Os04g0535200*, *Os08g0267300*, *Os05g0403000*, *Os04g0330900*, and *OsSAG12-1*) in endosperm (Fig. 7B).

In addition, this comparison method revealed that the influence of embryo on IAA was similar with T6P but converse with SnRK1 activity across the developmental stages, decreasing at 10 DAF while increasing at 15-25 DAF (Fig. 7A). Recently, by modulating T6P content in growing embryos of garden pea (*Pisum sativum*), Meitzel et al. (2021) found that auxin acts downstream of T6P to facilitate seed filling. Our finding of the synchronously dynamic pattern of IAA and T6P indicates a similar role of auxin-T6P pathway in grain-filling process of rice.

## DISCUSSION

Embryo-endosperm relationship is one of the determining factors contributing to seed development and subsequently the yield and quality of cereal grains. However, signals or regulators that coordinate the developmental processes of the two compartments remain largely unknown. This study took advantages of the NB mutant to evolve a novel comparison method for quantifying the influence of embryo on endosperm development, revealing that the embryo has a dragging effect on the developmental transition of the endosperm. To our knowledge, it is the first attempt to directly disclose the evidence of embryo-endosperm interaction during rice seed development. The findings reveal new aspects of the role of embryo in the formation process of rice quality, and help to draw an integrative picture of seed development from the perspective of agronomy and crop physiology.

### 1. The dragging effect of embryo on endosperm development

Seed development is an orchestrated progression through a series of stages, which can be described by two types of ‘time’: the chronological time and the developmental time defined by the sequence of stages (Ebisuya and Briscoe, 2018). There is evidence of bidirectional interaction between embryo and endosperm throughout development (An et al., 2020). Notably, two recent studies in Arabidopsis suggest an independent relationship between the two tissues. By analyzing mutants with defective endosperm cellularization, O’Neill et al. (2019) found that this endosperm process is not required for the onset of embryo maturation. Using single-fertilization mutants, Xiong et al. (2021) demonstrated that in the absence of embryo, endosperm develops the same way as in the wild type, suggesting that endosperm development is an autonomously programmed process independent of embryogenesis. These two studies in combination indicate that in terms of developmental time, the mechanisms controlling endosperm and embryo development act independently of each other (O’Neill et al., 2019). On the contrary, in terms of chronological time, the current study show that duration of endosperm filling in the bottom part of the NB grains was prolonged by the embryo (Fig. 2J), suggesting a substantial interaction between embryo and endosperm. Similarly, Xiong et al. (2021) showed that rapid embryo expansion can significantly accelerate endosperm breakdown, thus shortening the lifespan of the transient endosperm. Further, it should be noted that cereal seeds like rice have a persistent endosperm, whereas dicots like Arabidopsis have an ephemeral endosperm that degrades as the embryo grows. As a result, whether the independence of endosperm development from embryo in Arabidopsis is applicable to rice awaits further investigation.

Plants have evolved a variety of timing mechanisms that integrate chronological time with developmental time to ensure proper development (O’Neill et al., 2019). For the endosperm, these include internal timers of molecular oscillators based on hormones or metabolites, and external timers dependent on environmental signals or emanating from a different tissue like the embryo. This study reveals a dragging effect of embryo on endosperm development in chronological time, i.e. extending the storage accumulating stage whereas delaying the maturation stage. This finding provides direct evidence for the role of embryo as the external timer controlling endosperm development. In addition, hormones like GA, auxins, and ABA were unevenly distributed in rice seed, with the embryo generally having higher contents at the early and middle stages, as also reported by Zhang et al. (2020a). Interestingly, *GA20ox* (*GA20ox2* and *GA20ox3*), *GA3ox* (*OsGA3ox1* and *OsGA3ox2*), *KS* (*OsKS1* and *OsKS4*) were predominantly expressed in embryo at 5-20 DAF, indicating that embryo may be the GA-synthetic site whereas endosperm is the GA-acting site (Fig. 4F). It is well established that embryo-derived GA modulates the secretion of starch-degrading enzymes like α-amylase from the aleurone and scutellum upon germination. But it is still uncertain whether the degradation of starch during seed development is analogous to the germinating process (Sreenivasulu et al., 2015). Our results imply that one of the external timers coordinating rice grain development might be the hormone GA released from endosperm, which needs further investigation.

For internal timers modifying endosperm development, the T6P-SnRK1 signaling pathway may be the key component. At early stage of 5 to 10 DAF, sucrose in endosperm was lower, probably due to the deprivation by the growing embryo. In response to the reduced sucrose content, T6P was decreased simultaneously in endosperm, thus relieving the inhibition of SnRK1 activity. The increased SnRK1 activities promoted the catabolism or suppressed the anabolism of starch and proteins, as reflected by the lower content of starch and prolamins as well as the enhanced gene activity of amylase, lipase, and protease in endosperm (Fig. 7). Conversely, at middle stage between 20 and 25 DAF, the T6P-SnRK1 signaling showed an opposite trend relative to that between 5 and 10 DAF, indicating it may be involved in the transition of developmental stages in endosperm (Fig. 7). Taken together, the anticorrelation between T6P and SnRK1 activity opens the possibility that the T6P-SnRK1 pathway may be a master regulator coordinating the communication between embryo and endosperm during rice grain formation.

### 2. The integrative landscape of rice seed development

One of the major goals of crop production is to produce more grains with less time, thus improving resource use efficiency. The duration of each stage of seed development and the timing of transition between them is of agronomical significance. As reviewed by Olsen (2020), mechanism regulating the timing of endosperm cellularization has attracted attention due to its positive correlation with endosperm and seed size. Consequently, a precise and integrative landscape of seed development is necessary and useful for both fundamental and applied studies on the mechanisms underpinning crop yield and quality. Previously, from viewpoint of crop physiology, we proposed a practical staging system with three phases: embryo morphogenesis, endosperm filling, and seed maturation (An et al., 2020). Here, we update this staging system by integrating of molecular, physiological, and anatomical evidence in the current study as well as from the literature. In particular, considering the dragging effect of embryo on endosperm development at the former second stage (endosperm filling), we highlight the importance of this stage as critical for grain filling and quality formation, and thus subdivide it into two stages: embryo enlargement and endosperm filling. Accordingly, we paint a holistic and dynamic picture of rice seed development (Fig. 8), and provide a brief description of each stage and its agronomical relevance as follows.

**Fig. 8.**
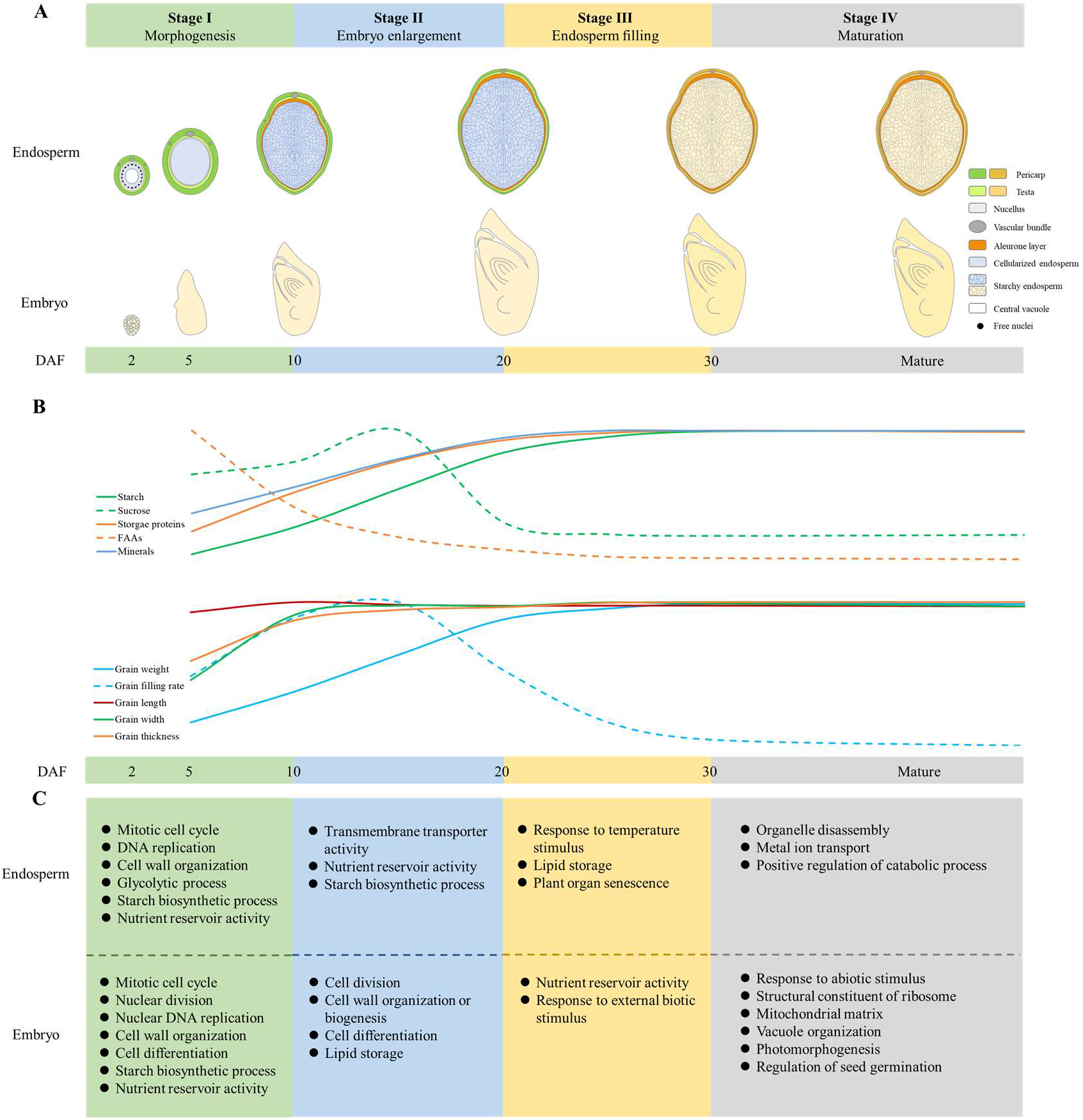
Holistic and dynamic picture of seed development. (A) Schematic illustration of the morphological changes in embryo (longitudinal section) and endosperm (transversal). Varying colors of the pericarp and testa show the progression of degradation in maternal tissues, while those of the starch endosperm show the grain-filling process. (B) Dynamic accumulation of storage materials (sucrose, starch, FAAs, storage proteins, and minerals) and the increase in grain weight, length, width, and thickness. (C) Molecular signatures of the embryo and endosperm at different developmental stages.

**Stage I**: Morphogenesis (0-10 DAF): After double fertilization, patterning and differentiation occurs simultaneously in the embryo and endosperm. At the end of this stage, most of morphogenetic events in embryo have already occurred. Endosperm has finished differentiation, forming two subregions, the aleurone and starchy endosperm, and starts storing starch and proteins.

**Stage II**: Embryo enlargement (10-20 DAF): Embryo greatly grows to its maximum volume at 20 DAF (Itoh et al., 2005). Starchy endosperm attains its highest rate of storage accumulation (Zhu et al., 2011; Fu et al., 2013), while the aleurone cells is filled with aleurone particles and spherosomes at the end of this stage (Yu et al., 2014). This stage witnesses strong interactions between embryo and endosperm, hence being critical for rice grain filling and quality.

**Stage III**: Endosperm filling (20-30 DAF): Embryo becomes dormant and endosperm continues to accumulate starch and proteins, reaching its maximum weight at 30 DAF. By the end of this stage, most of the starchy endosperm and maternal tissues have undergone PCD, losing their biological activity, while the aleurone and embryo is still alive (Wu et al., 2016b, c).

**Stage IV**: Maturation (30 DAF-maturity): After completion of reserve accumulation, the embryo becomes tolerant of desiccation, and undergoes a developmentally programmed dehydration event leading to dormancy and a quiescent state (Manfre et al., 2009; Angelovici et al., 2010). The starchy endosperm cells die completely upon seed maturation and desiccation. At this stage, seeds are susceptible to germination under hot and humid conditions, thus being vulnerable to preharvest sprouting (Du et al., 2018).

Fig. 8 draws distinctive patterns of embryo and endosperm development, with endosperm ceasing storage accumulation at 30 DAF, ten days after the corresponding timepoint of embryo (20 DAF). Supplementary Fig. S10 shows that the capacity of embryo to germinate starts at 15 DAF and peaks at 20 DAF for both genotypes of WT and NB. It is thus inferred that the embryo has the priority of nutrient allocation over endosperm during seed development. Moreover, it appears that this asynchrony of embryo and endosperm development is conserved across modern cultivars. They share a common chronological time, with embryo developmentally maturing at 20 DAF (Itoh et al., 2016; Armenta-Medina et al., 2021), while endosperm matures at 30 DAF (Morita et al., 2005; Yang et al., 2006; Wang et al., 2008; Chen et al., 2013; Wu et al., 2016c; Wei et al., 2017; Xu et al., 2021). Therefore, the essential period of rice yield and quality formation is the Stages I-III (Fig. 8), from anthesis to 30 DAF. After that, rice seed enters the stage of desiccation and maturation, lasting for 20 to 40 days with no marked increase in grain weight. By dividing the 60-day period of grain filling into two separate months, our previous report showed that only 10 % of the grain yield was formed after 30 DAF (Xu et al., 2021). Under the intensive rice-wheat and rice-oilseed rape cropping systems in the lower reaches of Yangtze River, China, late maturity of the rice crop causes the late sowing of wheat and oilseed rape, adversely affecting the seedling growth and subsequently the grain or seed yield (Bai and Tao, 2017; Zhang et al., 2020b).

Accordingly, we suggest a genetical intervention to shorten the duration of the late maturation stage of rice in order to adapt to the double cropping systems. In addition, future work should exploit more genotypes to verify if this asynchrony of embryo and endosperm development in chronological time is conserved in rice as well as other cereal crops like maize and wheat.

## CONCLUSIONS

Despite a number of comprehensive studies on the transcriptome atlas of seed development with high spatio-temporal resolutions, definite information about the interaction between the two economically important tissues of embryo and endosperm is still lacking. In this study, we successfully applied a novel comparison method based on the NB mutant, demonstrating a direct effect of embryo on the developmental processes of endosperm. Integrated analysis of transcriptome and metabolites identified putative regulatory timers that coordinate the developmental process between embryo and endosperm. The external timers for endosperm may be hormones like GA secreted from the embryo, while the internal timers be the T6P-SnRK1 signaling pathway that mediates carbon allocation in endosperm. In combination with results from molecular, physiological, and anatomical investigations, we framed a holistic and dynamic landscape of rice seed development, and explained its agronomical significance, in particular for the intensive systems of rice-wheat or rice-oilseed rape double cropping. In contrast to the quantum leap in fundamental science of genetic control of seed development, progress on the applied science like the physiology of grain filling and quality has apparently lagged behind. The integrative picture of rice grain formation proposed here is of value to bridge this knowledge gap, and will facilitate the designing of cogent strategies to enhance rice yield and quality.

## SUPPLEMENTARY DATA

The following supplementary data are available at *JXB* online.

Table S1. Details of primers used in this study.

Fig. S1. Correlation of gene expression level (FPKM) among biological replicates.

Fig. S2. qRT-PCR validation of RNA-Seq results.

Fig. S3. Expression profiles of tissue-specific genes in embryo and endosperm tissues.

Fig. S4. Morphology of the developing embryo and endosperm for WT and NB.

Fig. S5. GO functional categories enriched in different coexpression modules of embryo and endosperm for WT and NB, respectively.

Fig. S6. Carbohydrates, proteins, SnRK1, and hormones, and their related genes in the embryo and endosperm of NB throughout development.

Fig. S7. Fructose, glucose, and free amino acids in embryo and endosperm of WT.

Fig. S8. Fructose, glucose, and free amino acids in embryo and endosperm of NB.

Fig. S9. Minerals in the embryo and endosperm of NB throughout development.

Fig. S10. Germination rate of developing seed of WT and NB.

## ACKNOWLEDGEMENTS

The research was supported by the National Key R&D Program, Ministry of Science and Technology, China [2017YFD0300103], the National Natural Science Foundation of China (31771719), and National High Technology Research and Development Program of China (2014AA10A605). Rothamsted Research receives strategic funding from the Biotechnological and Biological Sciences Research Council of the UK. MP acknowledges funding from the Designing Future Wheat Strategic Programme (BB/P016855/1).

## AUTHOR CONTRIBUTIONS

YT had the main responsibility for data collection and analysis. YT and LA made the figures. FX, GL, YD and MP revised the manuscript. ZL had the overall responsibility for experimental design, project management and manuscript preparation. All authors contributed to the article and approved the submitted version.

## DATA AVAILABILITY STATEMENT

All the analysed data related to this manuscript are present in this paper and its supplementary data. All the raw data associated with this study are submitted to NCBI and are available with accession number PRJNA722833.

## REFERENCES

An L, Tao Y, Chen H, He MJ, Xiao F, Li GH, Ding YF, Liu ZH. 2020. Embryo-Endosperm Interaction and Its Agronomic Relevance to Rice Quality. Frontiers in Plant Science 11, 587641.

Angelovici R, Galili G, Fernie AR, Fait A. 2010. Seed desiccation: a bridge between maturation and germination. Trends in Plant Science 15, 211–218.

Armenta-Medina A, Gillmor CS, Gao P, Mora-Macias J, Kochian LV, Xiang D, Datla R. 2021. Developmental and genomic architecture of plant embryogenesis: from model plant to crops. Plant Communications 2, 100136.

Ashburner M, Ball CA, Blake JA, et al. 2000. Gene ontology: tool for the unification of biology. The Gene Ontology Consortium. Nature Genetics 25, 25–29.

Bai HZ, Tao FL. 2017. Sustainable intensification options to improve yield potential and eco-efficiency for rice-wheat rotation system in China. Field Crops Research 211, 89–105.

Belmonte MF, Kirkbride RC, Stone SL, et al. 2013. Comprehensive developmental profiles of gene activity in regions and subregions of the *Arabidopsis* seed. Proceedings of the National Academy of Sciences of the United States of America 110, E435–E444.

Bian JX, Deng PC, Zhan HS, Wu XT, Nishantha MDLC, Yan ZG, Du XH, Nie XJ, Song WN. 2019. Transcriptional Dynamics of Grain Development in Barley (*Hordeum vulgare* L.). International Journal of Molecular Sciences 20, 962.

Chen J, Zeng B, Zhang M, Xie SJ, Wang GK, Hauck A, Lai JS. 2014. Dynamic Transcriptome Landscape of Maize Embryo and Endosperm Development. Plant Physiology 166, 252–264.

Chen PF, Chen L, Jiang ZR, Wang GP, Wang SH, Ding YF. 2018a. Sucrose is involved in the regulation of iron deficiency responses in rice (*Oryza sativa* L.). Plant Cell Reports 37, 789–798.

Chen TT, Xu YJ, Wang JC, Wang ZQ, Yang JC, Zhang JH. 2013. Polyamines and ethylene interact in rice grains in response to soil drying during grain filling. Journal of Experimental Botany 64, 2523–2538.

Chen, Y.X., Chen, Y.S., Shi, C.M, et al. 2018b. SOAPnuke: a MapReduce acceleration-supported software for integrated quality control and preprocessing of high-throughput sequencing data. Gigascience 7,1–6.

Chou CC, Chen WS, Huang KL, Yu HC, Liao LJ. 2000. Changes in cytokinin levels of Phalaenopsis leaves at high temperature. Plant Physiology and Biochemistry 38, 309–314.

Delatte TL, Selman MHJ, Schluepmann H, Somsen GW, Smeekens SCM, de Jong GJ. 2009. Determination of trehalose-6-phosphate in *Arabidopsis* seedlings by successive extractions followed by anion exchange chromatography-mass spectrometry. Analytical Biochemistry 389, 12–17.

Dingkuhn M, Luquet D, Fabre D, Muller B, Yin XY, Paul MJ. 2020. The case for improving crop carbon sink strength or plasticity for a CO_2_-rich future. Current Opinion in Plant Biology 56, 259–272.

Doll NM, Just J, Brunaud V, et al. 2020. Transcriptomics at Maize Embryo/Endosperm Interfaces Identifies a Transcriptionally Distinct Endosperm Subdomain Adjacent to the Embryo Scutellum. Plant Cell 32, 833–852.

Du L, Xu F, Fang J, et al. 2018. Endosperm sugar accumulation caused by mutation of *PHS8/ISA1* leads to pre-harvest sprouting in rice. Plant Journal 95, 545–556.

Ebisuya M, Briscoe J. 2018. What does time mean in development? Development 145, dev164368.

Fu J, Xu YJ, Chen L, Yuan LM, Wang ZQ, Yang JC. 2013. Changes in Enzyme Activities Involved in Starch Synthesis and Hormone Concentrations in Superior and Inferior Spikelets and Their Association with Grain Filling of Super Rice. Rice Science. 20, 120–128.

Howe E, Holton K, Nair S, Schlauch D, Sinha R, Quackenbush J. 2010. MeV: Multi Experiment viewer. In Ochs MF, Casagrande JT. Davuluri RV, eds. Biomedical Informatics for Cancer Research. Boston: Springer US, 267–277.

Ingram GC. 2020. Family plot: the impact of the endosperm and other extra-embryonic seed tissues on angiosperm zygotic embryogenesis. F1000research 9.

Ishimaru T, Parween S, Saito Y, Shigemitsu T, Yamakawa H, Nakazono M, Masumura T, Nishizawa NK, Kondo M, Sreenivasulu N. 2019. Laser Microdissection-Based Tissue-Specific Transcriptome Analysis Reveals a Novel Regulatory Network of Genes Involved in Heat-Induced Grain Chalk in Rice Endosperm. Plant Cell Physiology 60, 626–642.

Itoh JI, Nonomura KI, Ikeda K, Yamaki S, Inukai Y, Yamagishi H, Kitano H, Nagato Y. 2005. Rice plant development: from zygote to spikelet. Plant and Cell Physiology 46, 23–47.

Itoh JI, Sato Y, Sato Y, Hibara KI, Shimizu-Sato S, Kobayashi H, Takehisa H, Sanguinet KA, Namiki N, Nagamura Y. 2016. Genome-wide analysis of spatiotemporal gene expression patterns during early embryogenesis in rice. Development 143, 1217–1227.

Juliano BO, Bechtel DB. 1985. The rice grain and its gross composition. In Juliano BO, ed. Rice: Chemistry and Technology. St. Paul: American Association of Cereal Chemistry, 17–57.

Kawakatsu T, Yamamoto MP, Hirose S, Yano M, Takaiwa F. 2008. Characterization of a new rice glutelin gene *GluD-1* expressed in the starchy endosperm. Journal of Experimental Botany 59, 4233–4245.

Kim D, Langmead B, Salzberg SL. 2015. HISAT: a fast spliced aligner with low memory requirements. Nature Methods 12, 357–360.

Lafon-Placette C, Köhler C. 2014. Embryo and endosperm, partners in seed development. Current Opinion in Plant Biology 17, 64–69.

Li B, Dewey CN. 2011. RSEM: accurate transcript quantification from RNA-Seq data with or without a reference genome. BMC Bioinformatics 12, 323.

Lin ZM, Wang ZX, Zhang XC, Liu ZH, Li GH, Wang SH, Ding YF. 2017. Complementary Proteome and Transcriptome Profiling in Developing Grains of a Notched-Belly Rice Mutant Reveals Key Pathways Involved in Chalkiness Formation. Plant and Cell Physiology. 58, 560–573.

Lin ZM, Zhang XC, Yang XY, Li GH, Tang S, Wang SH, Ding YF, Liu ZH. 2014. Proteomic analysis of proteins related to rice grain chalkiness using iTRAQ and a novel comparison system based on a notched-belly mutant with white-belly. BMC Plant Biology 14, 163.

Lin ZM, Zheng DY, Zhang XC, Wang ZX, Lei JC, Liu ZH, Li GH, Wang SH, Ding YF. 2016. Chalky part differs in chemical composition from translucent part of japonica rice grains as revealed by a notched-belly mutant with white-belly. Journal of the Science of Food and Agriculture. 96, 3937–3943.

Manfre AJ, LaHatte GA, Climer CR, Marcotte WR, Jr. (2009). Seed dehydration and the establishment of desiccation tolerance during seed maturation is altered in the *Arabidopsis thaliana* mutant *atem6-1*. Plant and Cell Physiology 50, 243–253.

Martínez-Barajas E, Delatte T, Schluepmann H, de Jong GJ, Somsen GW, Nunes C, Primavesi LF, Coello P, Mitchell RA, Paul MJ. 2011. Wheat grain development is characterized by remarkable trehalose 6-phosphate accumulation pregrain filling: tissue distribution and relationship to SNF1-related protein kinase1 activity. Plant Physiology 156, 373–381.

Meitzel T, Radchuk R, McAdam EL, et al. 2021. Trehalose 6-phosphate promotes seed filling by activating auxin biosynthesis. New Phytologist 229, 1553–1565.

Morita S, Yonemaru J, Takanashi J. 2005. Grain growth and endosperm cell size under high night temperatures in rice (*Oryza sativa* L.). Annals of Botany 95, 695–701.

Ning HF, Qiao JF, Liu ZH, Lin ZM, Li GH, Wang QS, Wang SH, Ding YF. 2010. Distribution of proteins and amino acids in milled and brown rice as affected by nitrogen fertilization and genotype. Journal of Cereal Science 52, 90–95.

O’Hara LE, Paul MJ, Wingler A. 2013. How do sugars regulate plant growth and development? New insight into the role of trehalose-6-phosphate. Molecular Plant 6, 261–274.

Olsen OA. 2020. The Modular Control of Cereal Endosperm Development. Trends in Plant Science 25, 279–290.

O’Neill JP, Colon KT, Jenik PD. 2019. The onset of embryo maturation in *Arabidopsis* is determined by its developmental stage and does not depend on endosperm cellularization. Plant Journal. 99, 286–301.

Paul MJ, Oszvald M, Jesus C, Rajulu C, Griffiths CA. 2017. Increasing crop yield and resilience with trehalose 6-phosphate: targeting a feast-famine mechanism in cereals for better source-sink optimization. Journal of Experimental Botany 68, 4455–4462.

Qin ZZ, Tang XH, Pan GZ, He MY. 1990. Changes of endogenous ABA levels in rice embryo and endosperm and association with development and germination. Journal of Integrative Plant Biology 32, 448–455. (in Chinese)

R Core Team. 2013. R: A Language and Environment for Statistical Computing. R Foundation for Statistical Computing, Vienna. http://www.R-project.org/.

Ram H, Singh A, Katoch M, et al. 2020. Dissecting the nutrient partitioning mechanism in rice grain using spatially resolved gene expression profiling. Journal of Experimental Botany.

Sastre Toraño J, Delatte TL, Schluepmann H, Smeekens SCM, de Jong GJ, Somsen GW. 2012. Determination of trehalose-6-phosphate in *Arabidopsis thaliana* seedlings by hydrophilic-interaction liquid chromatography–mass spectrometry. Analytical and Bioanalytical Chemistry 403, 1353–1360.

Sato Y, Hong SK, Tagiri A, Kitano H, Yamamoto N, Nagato Y, Matsuoka M. 1996. A rice homeobox gene, *OSH1*, is expressed before organ differentiation in a specific region during early embryogenesis. Proceedings of the National Academy of Sciences of the United States of America 93, 8117–8122.

Sreenivasulu N, Butardo VM, Jr, Misra G, Cuevas RP, Anacleto R, Kavi Kishor PB. (2015). Designing climate-resilient rice with ideal grain quality suited for high-temperature stress. Journal of Experimental Botany 66, 1737–1748.

Tong XH, Wang YF, Sun AQ, Bello BK, Ni S, Zhang J. 2018. *Notched Belly Grain 4*, a Novel Allele of *Dwarf 11*, Regulates Grain Shape and Seed Germination in Rice (*Oryza sativa* L.). International Journal of Molecular Sciences 19.

Wang E, Wang JJ, Zhu XD, et al. 2008. Control of rice grain-filling and yield by a gene with a potential signature of domestication. Nature Genetics 40, 1370–1374.

Wang LK, Feng ZX, Wang X, Wang XW, Zhang XG. 2010. DEGseq: an R package for identifying differentially expressed genes from RNA-Seq data. Bioinformatics 26, 136–138.

Wang ZX, Zhang FF, Xiao F, Tao Y, Liu ZH, Li GH, Wang SH, Ding YF. 2018. Contribution of mineral nutrients from source to sink organs in rice under different nitrogen fertilization. Plant Growth Regulation. 86, 159–167.

Wei XJ, Jiao GA, Lin HY, Sheng ZH, Shao G.N, Xie LH, Tang SQ, Xu QG, Hu PS. 2017. *GRAIN INCOMPLETE FILLING 2* regulates grain filling and starch synthesis during rice caryopsis development. Journal of Integrative Plant Biology 59, 134–153.

Wu C, Cui KH, Wang WC, Li Q, Fahad S, Hu QQ, Huang JL, Nie LX., Peng SB. 2016a. Heat-induced phytohormone changes are associated with disrupted early reproductive development and reduced yield in rice. Scientific Reports 6, 34978.

Wu TY, Müller M, Gruissem W, Bhullar NK. 2020. Genome Wide Analysis of the Transcriptional Profiles in Different Regions of the Developing Rice Grains. Rice 13, 62.

Wu XB, Liu JX, Li DQ, Liu CM. 2016b. Rice caryopsis development I: Dynamic changes in different cell layers. Journal of Integrative Plant Biology 58, 772–785.

Wu XB, Liu JX, Li DQ, Liu CM. 2016c. Rice caryopsis development II: Dynamic changes in the endosperm. Journal of Integrative Plant Biology 58, 786–798.

Xiang DQ, Quilichini TD, Liu ZY, et al. 2019. The Transcriptional Landscape of Polyploid Wheats and Their Diploid Ancestors during Embryogenesis and Grain Development. Plant Cell 31, 2888–2911.

Xiong HX, Wang W, Sun MX. 2021. Endosperm development is an autonomously programmed process independent of embryogenesis. Plant Cell.

Xu HF, Wang ZX, Xiao F, Yang L, Li GH, Ding YF, Paul MJ, Li WW, Liu ZH. 2021. Dynamics of dry matter accumulation in internodes indicates source and sink relations during grain-filling stage of japonica rice. Field Crops Research 263, 108009.

Xue LJ, Zhang JJ, Xue HW. 2012. Genome-wide analysis of the complex transcriptional networks of rice developing seeds. PLoS One 7, e31081.

Yamamoto MP, Onodera Y, Touno SM, Takaiwa F. 2006. Synergism between RPBF Dof and RISBZ1 bZIP activators in the regulation of rice seed expression genes. Plant Physiology 141, 1694–1707.

Yang JC, Zhang JH, Wang ZQ, Liu K, Wang P. 2006. Post-anthesis development of inferior and superior spikelets in rice in relation to abscisic acid and ethylene. Journal of Experimental Botany 57, 149–160.

Yi F, Gu W, Chen J, et al. 2019. High Temporal-Resolution Transcriptome Landscape of Early Maize Seed Development. Plant Cell 31, 974–992.

Yu XR, Zhou L, Xiong F, Wang Z. 2014. Structural and Histochemical Characterization of Developing Rice Caryopsis. Rice Science. 21, 142–149.

Zhang JJ, Xue HW. 2013. *OsLEC1*/*OsHAP3E* participates in the determination of meristem identity in both vegetative and reproductive developments of rice. Journal of Integrative Plant Biology 55, 232–249.

Zhang XF, Tong JH, Bai AN, Liu CM, Xiao LT, Xue HW. 2020a. Phytohormone dynamics in developing endosperm influence rice grain shape and quality. Journal of Integrative Plant Biology 62,1625–1637.

Zhang YH, Primavesi LF, Jhurreea D, Andralojc PJ, Mitchell RAC, Powers SJ, Schluepmann H, Delatte T, Wingler A, Paul MJ. 2009. Inhibition of SNF1-related protein kinase1 activity and regulation of metabolic pathways by trehalose-6-phosphate. Plant Physiology 149, 1860–1871.

Zhang Z, Cong RH, Ren T, Li H, Zhu Y, Lu JW. 2020b. Optimizing agronomic practices for closing rapeseed yield gaps under intensive cropping systems in China. Journal of Integrative Agriculture 19, 1241–1249.

Zhu GH, Ye NH, Yang JC, Peng XX, Zhang JH. 2011. Regulation of expression of starch synthesis genes by ethylene and ABA in relation to the development of rice inferior and superior spikelets. Journal of Experimental Botany 62, 3907–3916.

